# Intramuscular injection of nerve growth factor as a model of temporomandibular disorder: nature, time-course, and sex differences characterising the pain experience

**DOI:** 10.1101/2022.12.13.520244

**Authors:** SM Schabrun, E Si, SK Millard, AKI Chiang, S Chen, NS Chowdhury, DA Seminowicz

## Abstract

**Background:** Temporomandibular disorder (TMD) is a common condition that frequently transitions to chronic symptoms. Experimental pain models that mimic the symptoms of clinical TMD may be useful in understanding the mechanisms, and sex differences, present in this disorder. Here we aimed to comprehensively characterise the nature and time-course of pain, functional impairment and hyperalgesia induced by repeated intramuscular injection of nerve growth factor (NGF) into the masseter muscle, and to investigate sex differences in the NGF-induced pain experience.

**Methods:** 94 healthy individuals participated in a longitudinal observational study with 30-day follow-up. NGF was injected into the right masseter muscle on Day 0 and Day 2. Participants attended laboratory sessions to assess pain (Numerical Rating Scale; NRS), functional limitation (mouth opening distance, Jaw Functional Limitation Scale; JFLS) and mechanical sensitization (pressure pain thresholds; PPTs) on Days 0, 2 and 5 and completed twice daily electronic pain dairies from Day 0 to day 30.

**Results:** Peak pain averaged 2.0/10 (95 % CI: 1.6-2.4) at rest and 4.3/10 (95 % CI: 3.9-4.8) on chewing. Pain-free mouth opening distance reduced from 5.0 cm (95 % CI: 4.8-5.1 cm) on Day 0 to 3.7 cm (95 % CI: 3.5-3.9 cm) on Day 5. The greatest reduction in PPTs was observed over the masseter muscle. Females experienced higher pain, greater functional impairment, and greater sensitivity to mechanical stimuli than males.

**Conclusion:** Intramuscular injection of NGF is a useful model with which to explore the mechanisms, and sex differences, present in clinical TMD.

## Introduction

Temporomandibular disorder (TMD) is a common musculoskeletal pain condition that transitions to chronic symptoms in 49% of individuals^1,2^. TMD-myalgia is characterised by pain in the muscles of mastication exacerbated by jaw movement (with or without referral), and the presence of functional limitation such as reduced mouth opening^3^. Despite considerable research effort, the mechanisms that underpin the transition to chronic TMD remain uncertain. Experimental models that can induce prolonged pain in humans are important for the identification of mechanisms that may drive the development of chronic TMD.

Repeated intramuscular injection of nerve growth factor (NGF) is a prolonged human experimental pain model that induces progressively developing muscle pain, functional limitation and mechanical hyperalgesia lasting up to 21 days. NGF is a neurotrophic protein with a key role in development of the peripheral pain-signalling and sympathetic nervous systems (for review see^4^). In adulthood, NGF acts as a pain mediator interacting with nociceptors through a range of mechanisms including sensitisation of peripheral nociceptive terminals, altered nociceptor transcription and sprouting of nociceptors^4^. Elevated NGF levels are associated with chronic pain in animals^5^ and in humans with chronic clinical pain^6^, and anti-NGF antibodies prevent hyperalgesia in animal models^7^ and act as analgesics in patients with osteoarthritis^8^ and low back pain^9^. These characteristics of NGF suggest it may provide a clinically and mechanistically relevant model of the transition to chronic musculoskeletal pain.

Several studies have used intramuscular injection of NGF into the masseter muscle to mimic pain and dysfunction associated with TMD in otherwise healthy humans, showing the development of mechanical allodynia, hyperalgesia, pain on function and functional limitation^10–15^. However, these studies have included relatively small numbers of participants (ranging from n=14-30). Here we aimed to build on prior work using a large sample to comprehensively characterise the nature and time-course of pain, functional impairment and hyperalgesia induced by repeated intramuscular injection of NGF into the masseter muscle, and to investigate sex differences in the NGF-induced pain experience.

## Methods

### Design

A longitudinal experimental design was used to follow healthy individuals, injected with nerve growth factor into the right masseter muscle on two occasions, over the course of 30 days. This study represents an interim analysis of the PREDICT trial that aims to undertake analytical validation of an electroencephalography and transcranial magnetic stimulation derived cortical biomarker signature in a human pain model of TMD. The PREDICT trial is prospectively registered on ClinicalTrials.gov (NCT04241562) and the protocol published^16^. Ethical approval was obtained from the University of New South Wales (HC190206) and the University of Maryland Baltimore (HP-00085371).

### Participants

Ninety-four healthy individuals (41 females:53 males; aged 25 ± 7 years [mean±SD]) were recruited between November 2020 and May 2022 using flyers, mailing lists and social media posts. As this study aimed primarily to observe the time-course and nature of NGF-induced symptoms, it was not appropriate to calculate a traditional power based sample size. Previous studies have shown sex differences in experimentally induced-pain responses with sample sizes of 12 to 15 per group^17–19^, suggesting sufficient power to answer our aim of exploring sex differences. Participants were excluded based on the following criteria: (1) inability or refusal to provide written consent, (2) not aged between 18-44 years, (3) presence of an acute pain disorder, (4) history or presence of any chronic pain disorder including migraine, (5) history or presence of any other medical or psychiatric complaint, (6) use of opioids or illicit drugs in the past 3 months, (7) current smoker or using nicotine replacements, (8) pregnant or lactating women, and (9) contraindicated for transcranial magnetic stimulation (e.g. metal implants, epilepsy).

### Experimental protocol

Participants attended three laboratory visits on Days 0, 2 and 5. Each laboratory visit included completion of questionnaires (Muscle Soreness Scale, McGill Pain Questionnaire, Diagnostic Criteria for TMD, Jaw Functional Limitation Scale) and assessment of pain-free mouth opening and pressure pain thresholds (PPTs). Electroencephalography (EEG) and Transcranial Magnetic Stimulation (TMS) assessments, not reported here, were taken as part of the larger PREDICT study. Intramuscular injection of NGF was given into the right masseter muscle at the end of each laboratory session on days 0 and 2. Participants completed twice daily electronic pain dairies from day 0 to day 30. The experimental protocol is detailed in Figure 1.

**Figure 1.**
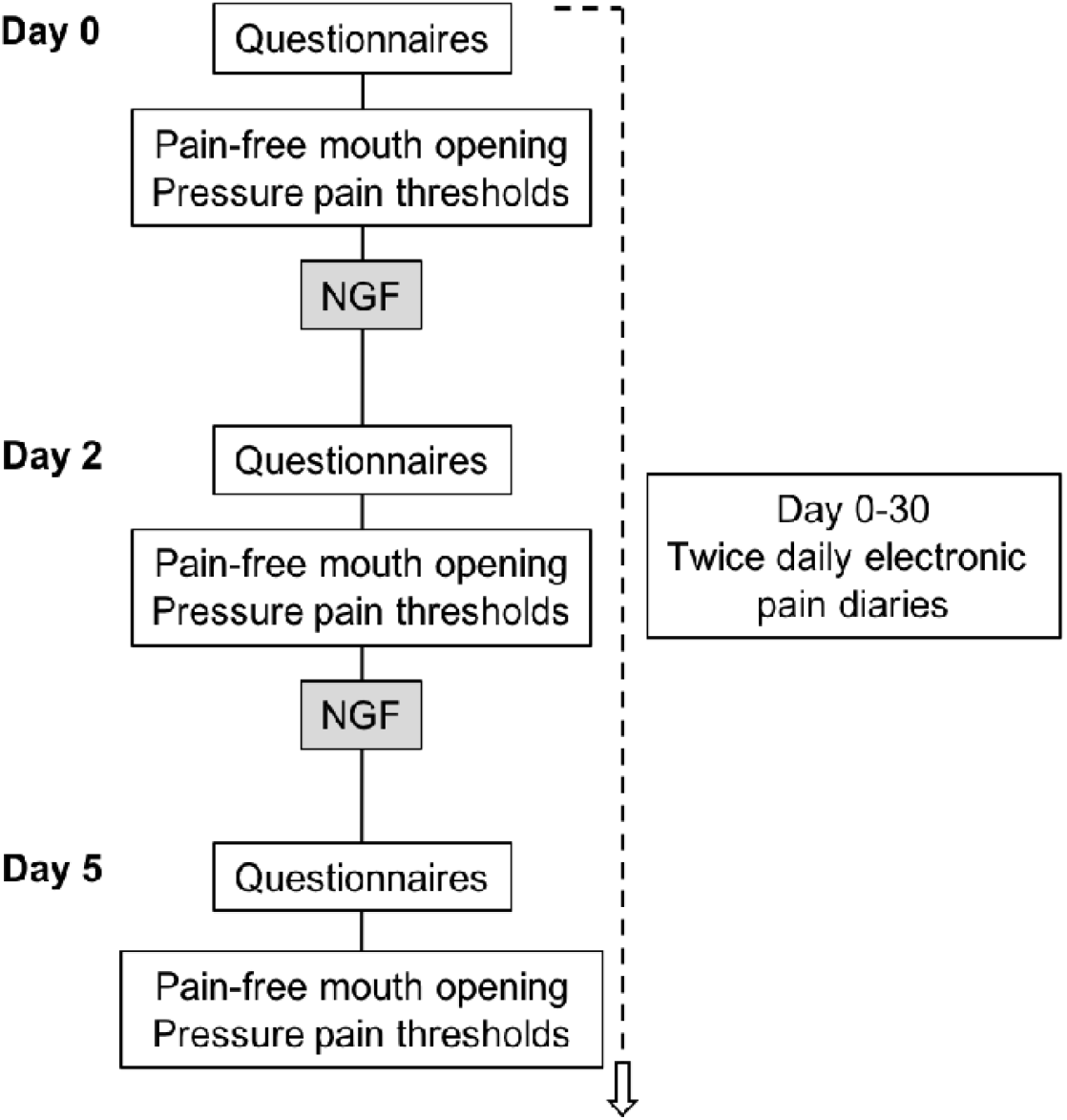
Experimental protocol

### Pain intensity and quality

Participants completed electronic pain diaries using a computer, tablet or phone at 10 am and 7 pm each day from day 0 to 30. Within the electronic diary, participants were asked to rate their pain intensity on an 11-point numerical rating scale anchored with ‘no pain’ at zero and ‘worst pain imaginable’ at 10 at rest and during chewing, swallowing, drinking, talking, yawning and smiling.

Muscle soreness was assessed using a modified 7-point Likert scale; 0 = “a complete absence of soreness,” 1 = “a light soreness in the muscle felt only when touched/vague ache,” 2 = “a moderate soreness felt only when touched/a slight persistent ache,” “3 = “a light muscle soreness when opening the mouth fully or chewing hard food,” 4 = “a light muscle soreness, stiffness or weakness when opening the mouth or chewing soft food,” 5 = “a moderate muscle soreness, stiffness or weakness when opening the mouth or chewing soft food,” and 6 = “a severe muscle soreness, stiffness or weakness that limits the ability to open the mouth or chew”.

The short-form McGill Pain Questionnaire^20^ was used to investigate the quality and distribution of NGF-induced muscle pain. Participants scored 15 pain descriptors (e.g. throbbing, aching, burning) as ‘none’, ‘mild’, ‘moderate’ or ‘severe’ on each day.

Participants were evaluated against the Diagnostic Criteria for TMD (DC-TMD)^3^ based on pain location, referral, palpation and the presence of headache to determine whether symptoms induced by NGF mimicked those associated with a clinical diagnosis of TMD.

### Functional impairment

Participants were asked to indicate the level of limitation experienced on a variety of functional tasks including chewing hard bread, opening the mouth wide enough to bite an apple, swallowing, talking etc. in the last 24 hours using the Jaw Functional Limitation Sale^21^. For each item, the participant rated the degree of functional impairment on an 11-point NRS anchored with ‘no limitation’ at zero and ‘severe limitation’ at 10. Items 1-6 represent mastication, items 7-10 mobility and items 11-20 verbal and emotional communication. Scores in each of these categories, as well as the total score, were used for analysis.

Pain-free mouth opening was assessed by asking participants to open their mouth as wide as possible without causing pain. Using a millimeter ruler, the distance between the incisal edge of the mandibular central incisor and the incisal edge of the opposing maxillary central incisor was recorded^22^.

### Pressure pain thresholds

Pressure pain thresholds were assessed at 5 sites: i) the right masseter muscle, ii) the right temporalis muscle, iii) the right temporomandibular joint, iv) the right trapezius muscle, and v) the right thumbnail. A 1-minute rest was observed between all measurements and 3 measures were made at each site. Pressure was applied at a rate of 40 kPa/s using a handheld algometer (Somedic, Horby, Sweden) perpendicular to the surface of the skin. Participants were asked to push a button when the sensation of pressure first changed to one of pain. The position of each site was recorded with a tape measure to ensure consistency of algometer placement across testing days. The average of the 3 trials at each site was used for analysis.

### Intramuscular injection of nerve growth factor

The injection site was sterilised with an alcohol wipe (70% Isopropyl, Bodichek®) and a sterile solution of recombinant human NGF (dose of 5 μg [0.2 mL]) given as a bolus injection into the right masseter muscle using a 1-mL syringe with a disposable needle (27 G). The needle was inserted perpendicular to the masseter body until bony contact was reached, retracted ∼2 mm, and NGF injected after a negative aspiration test^11^. Pain associated with NGF injection was reported using an 11-point numerical rating scale anchored with ‘no pain’ at zero and ‘worst pain imaginable’ at 10.

### Adverse events

Adverse events were monitored and reported according to National Institutes of Health guidelines. An adverse event was defined as an unexpected medical occurrence associated with the participant’s involvement in the study. The following details were reported: time of onset and offset, severity, likelihood of being related to the protocol, action taken and outcome.

### Data and statistical analyses

Data are reported using descriptive statistics (mean and 95% confidence intervals and percentages). Sex differences were compared for the following outcome variables: peak pain, pain duration, pain on injection, muscle soreness and total functional limitation score, using Mann-Whitney U tests. Bonferroni corrections were applied where appropriate.

Mouth opening distance and pressure pain threshold data were assessed for normality using the Shapiro-Wilk test and subsequently log-transformed. A 2-way mixed-model ANOVA with factors Sex (Male, Female) and Day (0, 2 and 5) was performed to assess changes in mouth opening distance with NGF injection and to explore a potential interaction between NGF-induced pain and sex.

A 3-way mixed-model ANOVA with factors Sex (Male, Female), Day (0, 2 and 5) and Muscle (masseter, temporalis, trapezius, TMJ, thumbnail) was performed to assess changes in pressure pain thresholds induced by NGF and to explore sex differences. To assess the change in PPT for each muscle, simple main effect contrasts were run comparing PPTs at Day 0 with Day 2 and Day 5 for each of the five muscle locations (Bonferroni corrected significance = .005). Additionally, two way-interaction contrasts compared these changes between males and females (Bonferroni corrected significance = .005). Finally, a planned two-way interaction contrast compared the magnitude of the change in PPTs in the masseter muscle against the magnitude of the change in PPTs for the remaining four muscles between i) Day 0 and Day 2 and ii) Day 0 and Day 5 (Bonferroni corrected significance = .025). A three-way interaction contrast compared these changes between males and females (Bonferroni corrected significance = .025).

## 3. Results

Nine participants had missing data. One participant withdrew from the study during the Day 0 testing session as they found the TMS procedure uncomfortable. Two participants were unable to attend the Day 2 test session due to illness and were excluded from analyses. Two participants were withdrawn from the study, and their data excluded from analyses, after they developed local swelling in the masseter muscle following the Day 0 injection. Four participants did not attend the Day 5 testing session (1 due to illness, 3 due to lockdown caused by the COVID-19 pandemic). These participants had missing data for pressure pain thresholds, pain-free mouth opening, diagnostic criteria, and all questionnaires for Day 5 but, as they had received both NGF injections and completed electronic pain diaries from day 1 to day 30, their data were retained for analyses. The percentage completion rate for the pain diaries across the 30-day period was 92%.

### 3.1 Pain intensity and quality

#### 3.1.1 Pain diary

Pain at rest and during chewing both peaked on the afternoon of Day 2 (Figure 2).

**Figure 2.**
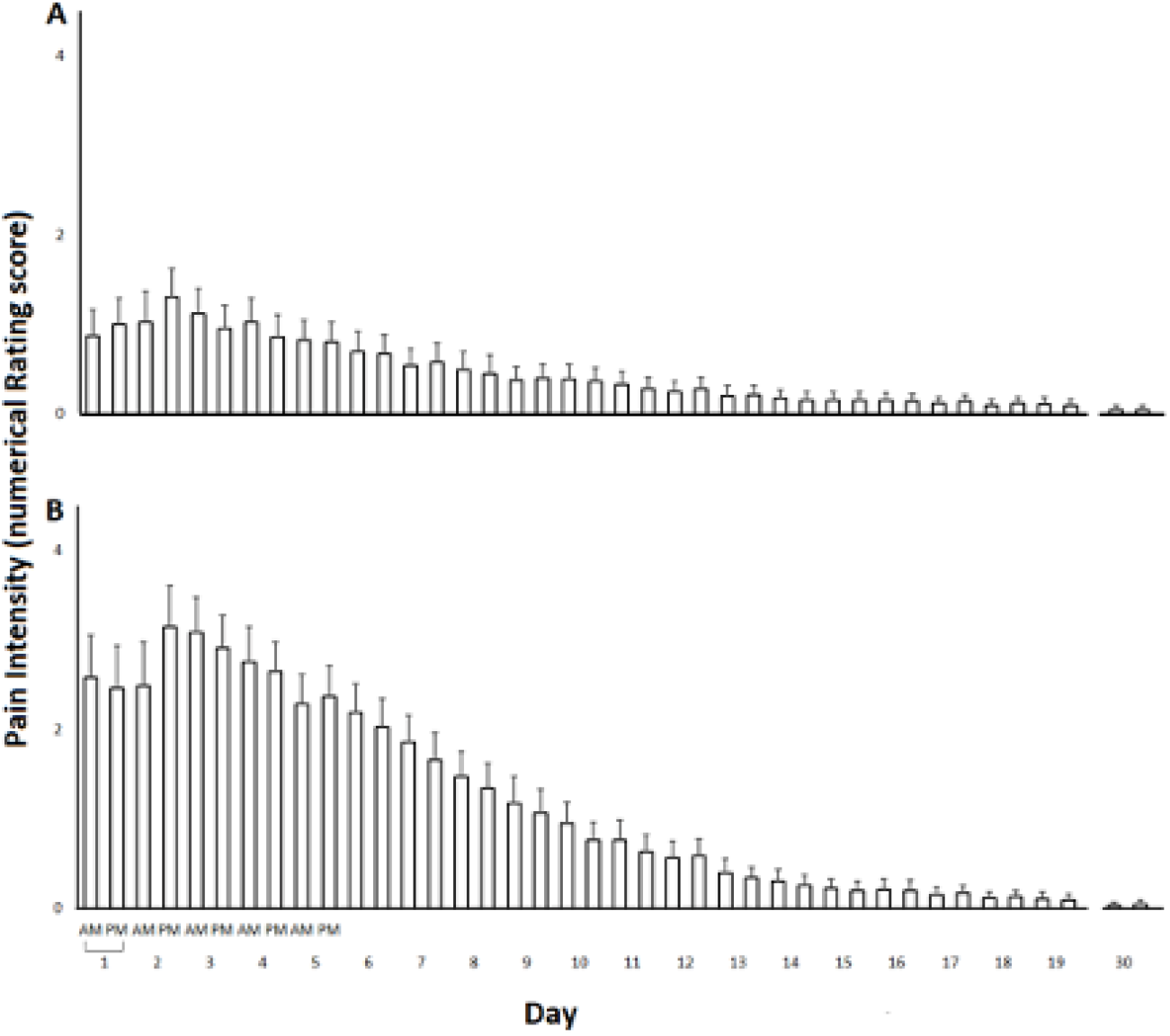
Intensity and time-course of nerve growth factor (NGF)-induced pain in the right masseter muscle (mean and 95 % CI) at rest (A) and during chewing (B). Pain was assessed using the 11-point numerical rating scale at 10am (AM) and 7pm (PM) each day from Day 1-30. NGF was injected on Day 0 and Day 2.

Average peak pain at rest was 2.0/10 (95 % CI: 1.6-2.4) while average peak pain during chewing was 4.3/10 (95 % CI: 3.9-4.8). Peak pain was significantly higher in females than in males (*U* = 671.50, *p* = .01) for pain on chewing, but not pain at rest (*U* =749, *p* = .052) (Figure 3A).

**Figure 3.**
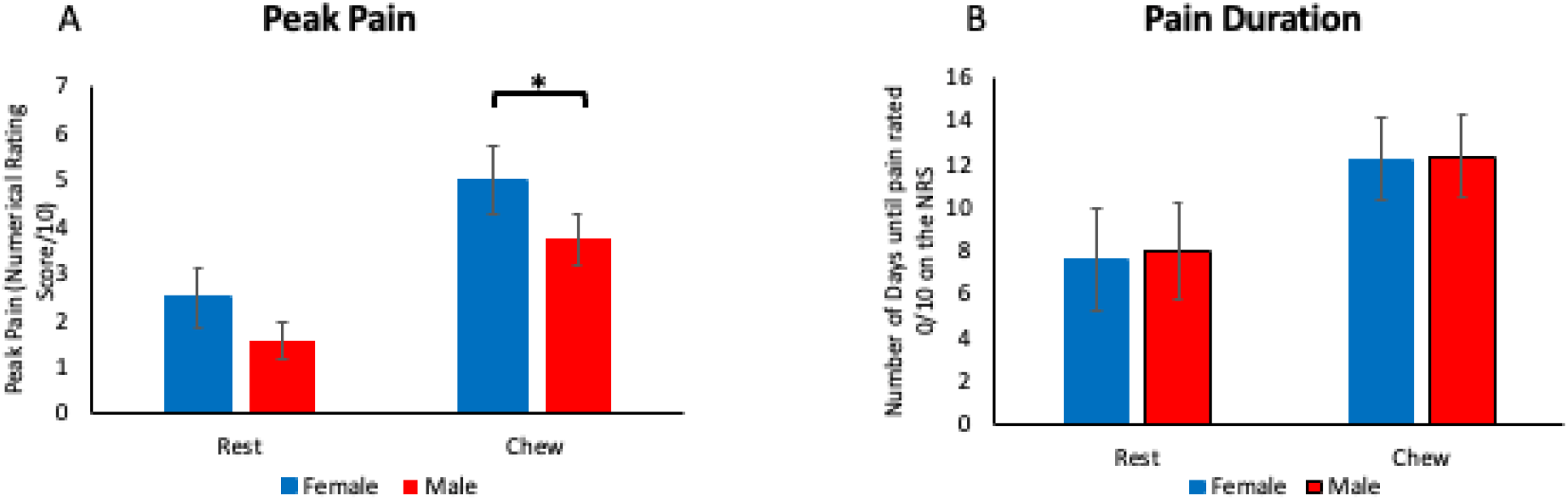
A: Peak pain intensity (mean and 95% CI) across the 30-day period at rest and during chewing, separated by sex. Peak pain was higher in females than in males for pain during chewing (* indicates p< .025). B: Duration of resting pain and pain on chewing (mean and 95% CI), separated by sex. Pain duration is defined as the number of days until pain was rated as 0/10 on the numerical rating scale (NRS). There were no differences in the duration of pain between males and females.

The average duration of pain was 7.9 days (95 % CI: 6.2 – 9.7) for pain at rest, and 12.5 days (95 % CI: 11.2-13.9) for pain on chewing. Pain had resolved completely in all but 4 participants by Day 25. Three participants provided ratings of 1/10 on the NRS at rest and during chewing from Day 25-Day 30. However, when contacted by the investigators for follow-up of unresolved pain at Day 30, all 3 participants reported that the 1/10 NRS scores were an error, and pain had in fact resolved prior to Day 25 (complete resolution of pain reported on Day 10, Day 17 and unable to recall exact day). The fourth participant continued to report mild pain local to the masseter muscle that did not resolve until Day 41. This was treated as an adverse event. There were no sex differences in pain duration (pain at rest: *U* =955.5, *p* = .84; pain on chewing: *U* =935.5, *p* =.71) (Figure 3B).

#### 3.1.2 McGill Pain Questionnaire

Pain quality was most frequently described as tender and aching on both Day 2 and 5. The frequency distribution was similar across Day 2 and 5 (Figure 4).

**Figure 4.**
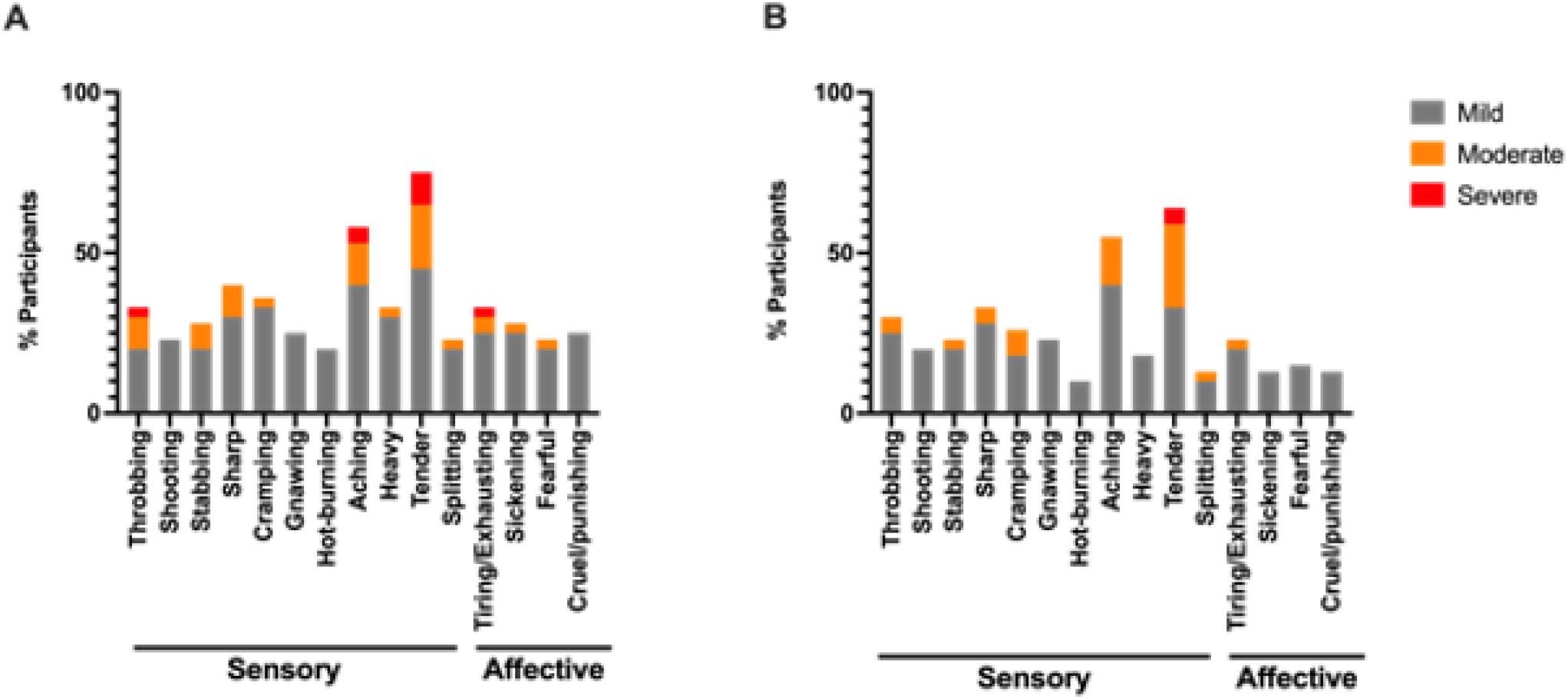
Percentage of participants reporting each category descriptor of pain on the McGill Pain Questionnaire on Day 2 (A) and Day 5 (B).

#### 3.1.3 Diagnostic criteria for TMD

Figure 5 shows the number of participants demonstrating symptoms in each diagnostic category of the DC-TMD on Day 0, 2, and 5. 84 participants (98%) reported symptoms consistent with the diagnostic criteria for local myalgia, myofascial TMD or myofascial TMD with referral, on both Day 2 and Day 5.

**Figure 5.**
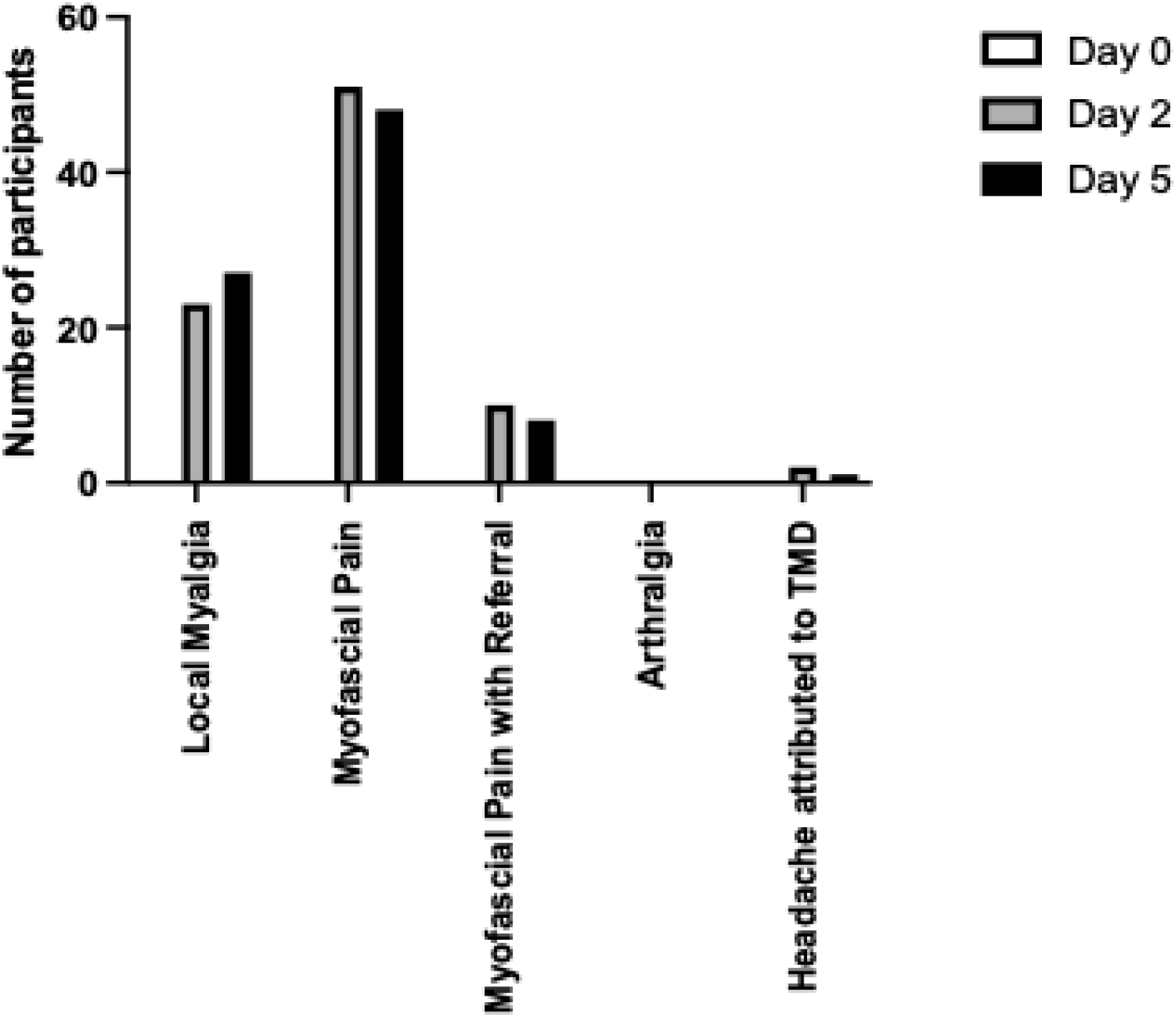
Number of participants classified in each of the DC-TMD categories on each testing day.

#### 3.1.4 Pain on injection

Pain on injection was 7.6 (95% CI: 7.2-7.9) out of 10 on Day 0 and 7.1 (95% CI: 6.7-7.5) out of 10 on Day 2. Pain associated with injection resolved rapidly, returning to baseline within a few minutes in all participants. Females reported higher pain on injection than males on Day 0 (*U* =579.5, *p* = .01), but not Day 2 (*U* =621.5, *p* =.18).

#### 3.1.5 Muscle soreness

Muscle soreness induced by NGF injection was reported on the 7-point Likert scale (mean and interquartile range) as 2.7 (2-3.8) on Day 2 and 2.9 (2-4) on Day 5, consistent with the category descriptor: a moderate soreness felt only when touched/a slight persistent ache. 15.2 % of individuals on Day 2 and 16.3 % of individuals on Day 5 reported moderate or severe muscle soreness (category 6 or 7). The frequency distribution of muscle soreness in each category, on each day, is presented in Table 1. Females reported greater levels of muscle soreness than males on Day 2 (*U* =495, *p* <.001), but not Day 5 (*U* =778, *p* =.28) (Figure 6A).

**Table 1:** Percent of participants reporting each category descriptor on the muscle soreness scale.

| Category | Day 0 | Day 2 | Day 5 |
| --- | --- | --- | --- |
| A complete absence of soreness | 100 | 0 | 0 |
| A light soreness felt only when touched/vague ache | 0 | 24.4 | 20.9 |
| A moderate soreness felt only when touched/a slight persistent ache | 0 | 25.6 | 22.1 |
| A light soreness when opening the mouth fully or chewing hard food | 0 | 24.4 | 20.9 |
| A light soreness, stiffness or weakness when opening the mouth or chewing soft food | 0 | 10.4 | 19.8 |
| A moderate soreness, stiffness or weakness when opening the mouth or chewing soft food | 0 | 14 | 15.1 |
| A severe soreness, stiffness or weakness that limits the ability to open the mouth or chew | 0 | 1.2 | 1.2 |

**Figure 6.**
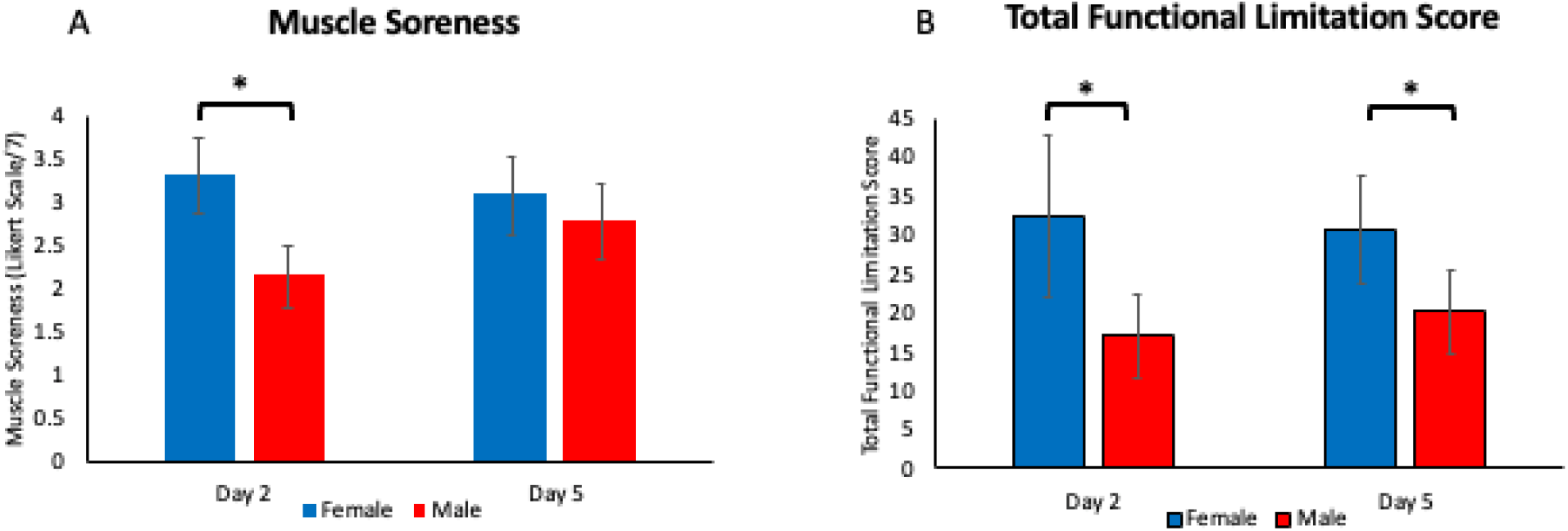
A: Muscle soreness score (mean and 95% CI) on Day 2 and Day 5, separated by sex. Muscle soreness was higher in females than in males on Day 2 (* indicates p< .025). B: Total functional impairment score (mean and 95% CI) on Day 2 and Day 5, separated by sex. Functional impairment was greater in females than in males on both Day 2 and Day 5 (* indicates p< .025).

### 3.2 Functional impairment

Total functional impairment scores were 24.1 (95 % CI: 18.0-30.2) on Day 2 and 25.0 (95 % CI: 20.4-29.6) on Day 5. The greatest functional impairment was observed in the mastication subscale (Table 2). Females reported greater functional impairment in response to NGF injection than males on both Day 2 (*U* =572.5, *p* = .004), and Day 5 (*U* =598, *p* =.01) (Figure 6B).

**Table 2.**
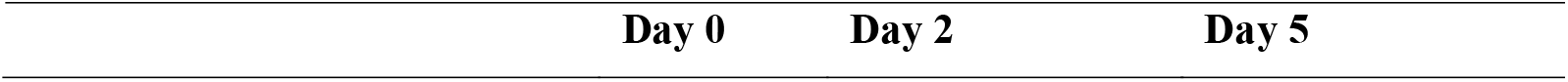
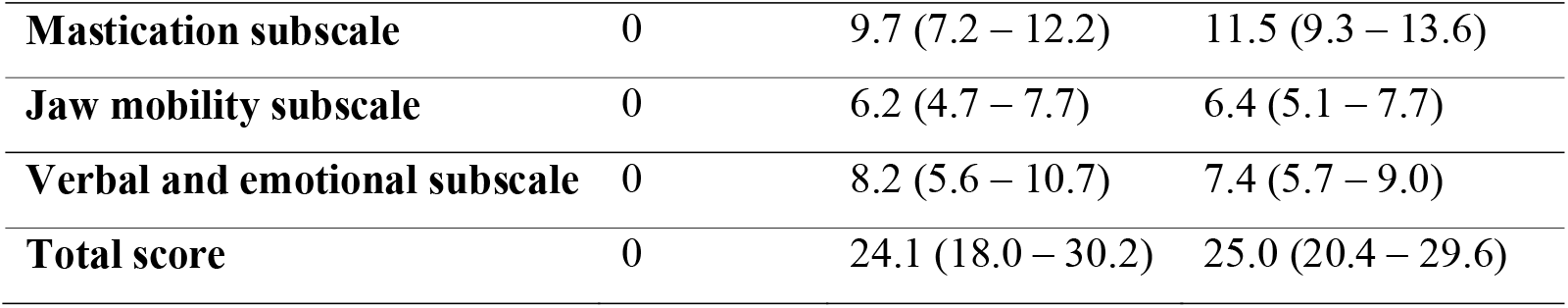
Jaw Functional Limitation Scale scores (mean and 95 % confidence interval)

| Day 0 | Day 2 | Day 5 |
| --- | --- | --- |

|  |  |  |  |
| --- | --- | --- | --- |
| **Mastication subscale** | 0 | 9.7 (7.2 – 12.2) | 11.5 (9.3 – 13.6) |
| **Jaw mobility subscale** | 0 | 6.2 (4.7 – 7.7) | 6.4 (5.1 – 7.7) |
| **Verbal and emotional subscale** | 0 | 8.2 (5.6 – 10.7) | 7.4 (5.7 – 9.0) |
| **Total score** | 0 | 24.1 (18.0 – 30.2) | 25.0 (20.4 – 29.6) |

Pain-free mouth opening distance reduced from 5.0 cm (95 % CI: 4.8-5.1 cm) on Day 0, to 4.0 cm (95 % CI: 3.7-4.2 cm) on Day 2 and 3.7 cm (95 % CI: 3.5-3.9 cm) on Day 5 (main effect of time: *F*_2, 162_ = 62.04, *p* < 0.001; time x sex interaction: *F*_2, 162_ = 76, *p* = 0.47). Females reported greater limitation to mouth opening distance over time than males (*F*_1, 81_ = 14.76, *p* < 0.001).

### 3.3 Pressure pain thresholds

Analyses of PPT data revealed a significant main effect of time (*F*_1.82, 151.13_ = 172.2, *p* < 0.001), an interaction effect between Day and Muscle *(F*_2,89, 239.65_ = 232.8, *p* < 0.001), and a three-way interaction effect between Day, Muscle and Sex (*F*_2,89, 239.65_ = 11.6, *p* < 0.001) (Figure 7).

**Figure 7.**
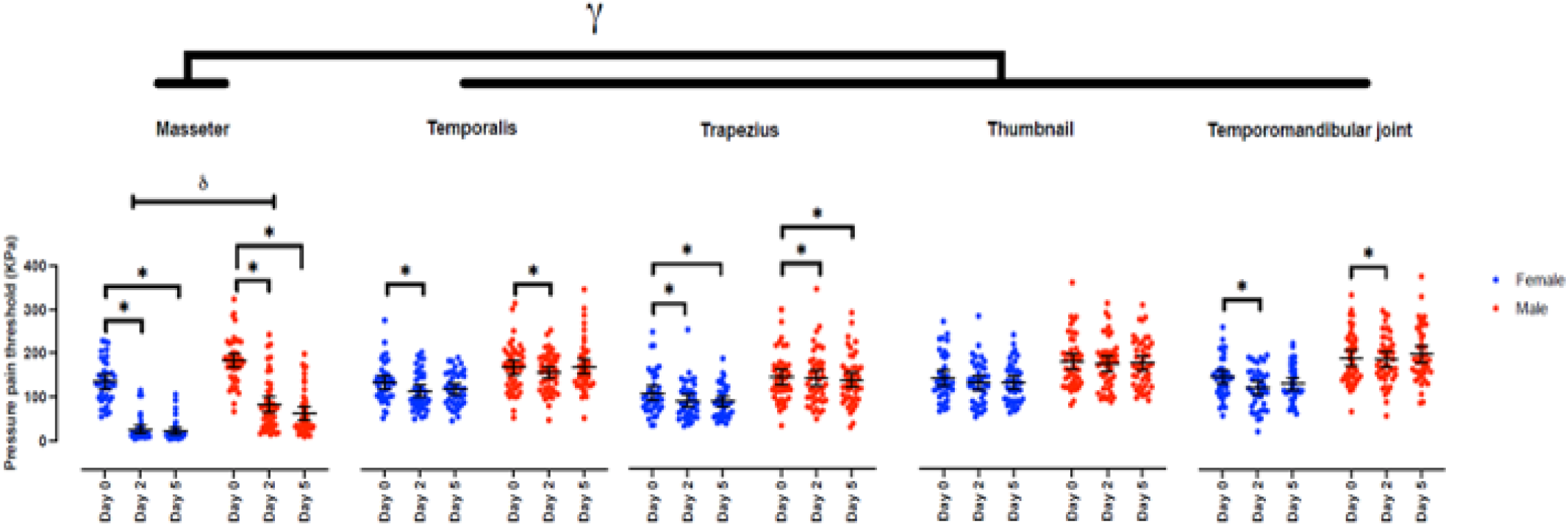
Pressure pain thresholds (PPTs; mean and 95 % CI) for each site on each testing day. Relative to Day 0, PPTs were reduced over the masseter muscle on Day 2 and Day 5 (* indicates p<0.005). PPTs were also reduced over the temporalis and TMJ sites on Day 2 (* indicates p<0.005), and over trapezius on Day 2 and Day 5 (* indicates p<0.005). The magnitude of the reduction in the PPTs was greater for the masseter muscle than for any other testing site on both Day 2 and 5 (□ indicates p<0.025). The magnitude of the reduction in the PPTs over the masseter muscle was larger in females than males on Day 2 and Day 5 (δ indicates p < 0.005).

Relative to Day 0, PPTs were reduced in the masseter muscle (*F*_1, 83_ =376.3, *p* < 0.001, *d* = 1.9) and the TMJ (*F*_1, 83_ =13.0, p = .002, *d* = 0.28) on Day 2. The magnitude of these reductions was greater in females than males (masseter: *F*_1, 83_ =33.8, *p* < 0.001, TMJ: *F*_1, 83_ =9.8, *p* = 0.002). PPTs were also reduced in the temporalis muscle (*F*_1, 83_ =20.6, *p* < 0.001, *d* = 0.31) and the trapezius muscle (*F*_1, 83_ =11.3, *p* = 0.001, *d* = 0.21) on Day 2. No sex differences were observed for these muscles (temporalis: *F*_1, 83_ = 4.3, *p* = .042, trapezius: *F*_1, 83_ = 5.0, *p* = .027).

Relative to Day 0, PPTs were reduced in the masseter muscle on Day 5 (*F*_1, 83_ =451.9, *p* < 0.001, *d* = 2.4), and the magnitude of the reduction was greater in females than males (*F*_1, 83_ =17.2, *p* < 0.001). PPTs were also reduced in the trapezius muscle on Day 5 (*F*_1, 83_ = 12.3, *p* = 0.001, *d* = 0.21), with no sex differences observed (*F*_1, 83_ = 3.0, *p* = .087).

The reduction in PPTs on Day 2 (*F*_1, 83_ =358.94, *p* < 0.001) and Day 5 (*F*_1, 84_ =418.82, *p* < 0.001) was greater for the masseter muscle than for any other site. Females reported a greater reduction in PPTs in the masseter (relative to other muscles) than males on both Day 2 (*F*_1, 83_ =37.3, *p* < 0.001) and Day 5 (*F*_1, 83_ =20.6, *p* < 0.001).

### 3.4 Adverse events

Three adverse events occurred. One participant reported local, soft swelling in both cheeks following the second NGF injection attributed to a diagnosed salivary gland condition that was not disclosed to the research team prior to study enrolment. All symptoms resolved within 48 hours. A second participant reported local, soft swelling of the right masseter muscle following the first NGF injection that resolved within 48 hours. A third participant reported pain in the right masseter muscle that lasted beyond the 30-day time-period. Pain induced by NGF injection in this participant was of average intensity (max pain of 3/10 reported on chewing on Day 2) and declined steadily throughout the study period (pain reduced to 1/10 on chewing on from Day 10 onwards) consistent with the expected NGF trajectory. Pain had resolved completely by Day 41.

## Discussion

This observational study of a large sample demonstrates that repeated intramuscular injection of NGF into the masseter muscle is safe and well tolerated. Two 5μg doses of recombinant human NGF, injected 48 hours apart, produced pain, functional impairment and mechanical hyperalgesia that mimicked the clinical characteristics of TMD, providing support for the use of NGF as a model to explore the neurophysiological mechanisms underpinning the human TMD pain experience. In addition, this study demonstrated sex differences in pain behaviour in response to NGF injection with females reporting higher peak pain on chewing, worse pain on initial NGF injection, greater muscle soreness, functional impairment and restriction to mouth opening distance, and greater sensitivity to mechanical stimuli than males. These findings add to a growing body of literature demonstrating sex differences in experimental and clinical pain populations.

A number of experimental pain models have been used to investigate the neurophysiological mechanisms contributing to musculoskeletal pain disorders, including TMD. However, these models are limited by the transient nature, and quality, of induced pain which is often inconsistent with that experienced by those with clinical pain. In the current study, two intramuscular injections of NGF into the masseter muscle produced symptoms consistent with the diagnostic criteria for local myalgia or myofascial TMD (with or without referral) in 98 % of participants. Participants reported pain predominantly localised to the masseter muscle that was of a tender and aching nature, was present at low intensities at rest and was exacerbated by oral function (e.g. chewing, yawning). Pain resulted in impaired function, with pain-free mouth opening distance reducing from 5.0 cm (95 % CI: 4.8-5.1 cm) on Day 0 to 3.7 cm (95 % CI: 3.5-3.9 cm) on Day 5. These values are consistent with pain free mouth opening distances reported in clinical TMD cohorts (e.g. chronic TMD 3.4±0.09 cm [mean±SEM]; healthy non-TMD individuals 4.7±0.04 cm)^23^. The level of disability reported on the Jaw Functional Limitation Scale was also similar between our NGF cohort and those with chronic clinical TMD^23^. Finally, sensitivity to pressure stimuli has been reported amongst individuals with clinical TMD. When pressure pain thresholds were assessed over the masseter muscle, non-TMD individuals displayed an average PPT of 195.5± 3.5 kPa while individuals with chronic TMD had an average PPT of 128.5± 4.6 kPa. Here we report PPTs of 161± 11.98 kPa before pain onset and 44± 9.59 kPa on Day 5, suggesting intramuscular injection of NGF induces greater mechanical sensitivity in otherwise healthy individuals than is observed in chronic TMD. This finding is not surprising given that NGF induces pain, at least in part, through sensitisation of high threshold mechanosensitive afferents^24^.

Previous studies in animals^25^ and humans^19^ have shown sex differences in the NGF-induced pain response. For example, the magnitude and duration of mechanical sensitivity induced by NGF injection into the masseter muscle is greater in female than male, rats^26^. Similarly, in humans, studies have shown significantly greater sensitization to mechanical stimuli, and greater pain during oral function in females than males^19^. Our findings confirm these sex differences in a large sample size. Of note, although mechanical sensitization was present for the masseter, temporalis, trapezius and TMJ recording sites in the current study, sex differences were only observed for the masseter and TMJ, suggesting some degree of specificity to the injected muscle in the sex response to NGF. Although the mechanism underpinning these sex differences remains unclear, previous studies have suggested increased expression of NMDA-receptors by masticatory muscle nerves in females following NGF injection that is associated with the degree of mechanical sensitization^19^. As women are known to experience clinical TMD more frequently than men^27,28^, intramuscular injection of NGF into the masseter muscle may be a useful model to further explore the mechanisms underpinning sex differences in the pain experience.

The present study administered intramuscular injection of NGF to the masseter muscle of 94 otherwise healthy individuals with few adverse events. Local swelling of the muscle is a known potential adverse event associated with intramuscular injection. In our study two incidents of localised muscle swelling were reported, one associated with the injection, and one with an undisclosed prior salivary gland condition. In both cases swelling resolved within 48 hours. However, we recommend future studies include specific screening for salivary gland conditions and exclude these individuals from participation. The final adverse event involved an expectedly long duration of low intensity muscle pain (1/10 on the NRS from Day 10 to Day 41). The reason for this extended duration of pain is unclear. NGF-induced pain was of average intensity and there were no other features of the pain experience that stood-out as potentially predisposing this individual to a longer pain duration. The extended follow-up period in the current study is a key strength of our protocol allowing detailed observation of the resolution stage of NGF-induced pain. Future studies should include sufficiently long follow-up periods to monitor individuals who experience longer pain durations and ensure all individuals experience complete resolution of pain before study exit.

This study was intended as a detailed observation of pain, function and mechanical sensitivity induced by NGF injection into the human masseter muscle. To our knowledge, the twice daily assessments of pain for 30 days in a large sample of 53 males and 41 females represents the most in-depth examination of the nature and time-course of NGF-induced pain in the current literature. However, some limitations exist. For example, tests of pressure pain thresholds and mouth-opening distance were only performed on Days 0, 2 and 5 limiting insight into the time-course of these features of the NGF pain experience. Future studies should seek to assess these variables across the 30-day follow-up period. In addition, this study does not include examination of mechanisms that could explain sex differences in the pain experience. More studies investigating a variety of biopsychosocial mechanisms are needed to increase our understanding of why females experience greater pain, greater functional limitation and higher mechanical sensitivity than males.

## Conclusion

Repeated intramuscular injection of NGF into the masseter muscle induces pain and functional limitation similar to that of clinical TMD. Females experience higher pain on oral function, greater functional impairment, and greater sensitivity to mechanical stimuli than males. The NGF pain model may be useful as a tool to further understand the mechanisms, and sex differences, involved in clinical TMD.

## Notes

**Disclosures** This project was funded by grant 1R61NS113269-01 from The National Institutes of Health to DAS, SMS and SC. DAS is an advisor to Empower Therapeutics. The other authors have no conflicts to declare.

### Competing Interest Statement

This project was funded by grant 1R61NS113269-01 from The National Institutes of Health to DAS, SMS and SC. DAS is an advisor to Empower Therapeutics. The other authors have no conflicts to declare.

## References

1. Slade, G.D., et al. Signs and symptoms of first-onset TMD and sociodemographic predictors of its development: the OPPERA prospective cohort study. J Pain 14, T20–32 e21-23 (2013).

2. Slade, G.D., et al. Painful Temporomandibular Disorder: Decade of Discovery from OPPERA Studies. J Dent Res 95, 1084–1092 (2016).

3. Schiffman, E., et al. Diagnostic Criteria for Temporomandibular Disorders (DC/TMD) for Clinical and Research Applications: recommendations of the International RDC/TMD Consortium Network* and Orofacial Pain Special Interest Groupdagger. J Oral Facial Pain Headache 28, 6–27 (2014).

4. Denk, F., Bennett, D.L. & McMahon, S.B. Nerve Growth Factor and Pain Mechanisms. Annu Rev Neurosci 40, 307–325 (2017).

5. Isola, M., et al. Nerve growth factor concentrations in the synovial fluid from healthy dogs and dogs with secondary osteoarthritis. Vet Comp Orthop Traumatol 24, 279–284 (2011).

6. Halliday, D.A., Zettler, C., Rush, R.A., Scicchitano, R. & McNeil, J.D. Elevated nerve growth factor levels in the synovial fluid of patients with inflammatory joint disease. Neurochem Res 23, 919–922 (1998).

7. Koltzenburg, M., Bennett, D.L., Shelton, D.L. & McMahon, S.B. Neutralization of endogenous NGF prevents the sensitization of nociceptors supplying inflamed skin. Eur J Neurosci 11, 1698–1704 (1999).

8. Lane, N.E., et al. Tanezumab for the treatment of pain from osteoarthritis of the knee. N Engl J Med 363, 1521–1531 (2010).

9. Katz, N., et al. Efficacy and safety of tanezumab in the treatment of chronic low back pain. Pain 152, 2248–2258 (2011).

10. Svensson, P., Cairns, B.E., Wang, K. & Arendt-Nielsen, L. Injection of nerve growth factor into human masseter muscle evokes long-lasting mechanical allodynia and hyperalgesia. Pain 104, 241–247 (2003).

11. Costa, Y.M., et al. Masseter corticomotor excitability is decreased after intramuscular administration of nerve growth factor. Eur J Pain 23, 1619–1630 (2019).

12. Costa, Y.M., et al. Local anaesthesia decreases nerve growth factor induced masseter hyperalgesia. Sci Rep 10, 15458 (2020).

13. Svensson, P., Castrillon, E. & Cairns, B.E. Nerve growth factor-evoked masseter muscle sensitization and perturbation of jaw motor function in healthy women. J Orofac Pain 22, 340–348 (2008).

14. Svensson, P., Wang, K., Arendt-Nielsen, L. & Cairns, B.E. Effects of NGF-induced muscle sensitization on proprioception and nociception. Exp Brain Res 189, 1–10 (2008).

15. Zhang, Y., et al. Comparison of Pain-Generated Functional Outcomes in Experimental Models of Delayed-Onset Muscle Soreness and Nerve Growth Factor Injection of the Masticatory Muscles. J Oral Facial Pain Headache 34, 311–322 (2020).

16. Seminowicz, D.A., et al. A novel cortical biomarker signature for predicting pain sensitivity: protocol for the PREDICT longitudinal analytical validation study. Pain Rep 5, e833 (2020).

17. Svensson, P., et al. Glutamate-evoked pain and mechanical allodynia in the human masseter muscle. Pain 101, 221–227 (2003).

18. Cairns, B.E., Hu, J.W., Arendt-Nielsen, L., Sessle, B.J. & Svensson, P. Sex-related differences in human pain and rat afferent discharge evoked by injection of glutamate into the masseter muscle. J Neurophysiol 86, 782–791 (2001).

19. Alhilou, A.M., et al. Sex-related differences in response to masseteric injections of glutamate and nerve growth factor in healthy human participants. Sci Rep 11, 13873 (2021).

20. Melzack, R. The short-form Mcgill pain questionnaire Pain 30, 191–197 (1987).

21. Ohrbach, R., Larsson, P. & List, T. The jaw functional limitation scale: development, reliability, and validity of 8-item and 20-item versions. J Orofac Pain 22, 219–230 (2008).

22. Rauch, A. & Schierz, O. Reliability of mandibular movement assessments depending on TMD. Cranio 36, 156–160 (2018).

23. Fillingim, R.B., et al. Long-term changes in biopsychosocial characteristics related to temporomandibular disorder: findings from the OPPERA study. Pain 159, 2403–2413 (2018).

24. Hoheisel, U., Unger, T. & Mense, S. Excitatory and modulatory effects of inflammatory cytokines and neurotrophins on mechanosensitive group IV muscle afferents in the rat. Pain 114, 168–176 (2005).

25. Syrett, M., et al. Sex-Related Pain Behavioral Differences following Unilateral NGF Injections in a Rat Model of Low Back Pain. Biology (Basel) 11(2022).

26. Wong, H., et al. NGF-induced mechanical sensitization of the masseter muscle is mediated through peripheral NMDA receptors. Neuroscience 269, 232–244 (2014).

27. LeResche, L. Epidemiology of temporomandibular disorders: implications for the investigation of etiologic factors. Crit Rev Oral Biol Med 8, 291–305 (1997).

28. Isong, U., Gansky, S.A. & Plesh, O. Temporomandibular joint and muscle disorder-type pain in U.S. adults: the National Health Interview Survey. J Orofac Pain 22, 317–322 (2008).

